# Pool walking may temporarily improve renal function by suppressing renin-angiotensin-aldosterone system in pregnant women

**DOI:** 10.1101/553081

**Authors:** Tatsuya Yoshihara, Masayoshi Zaitsu, Shiro Kubota, Hisatomi Arima, Toshiyuki Sasaguri

**Affiliations:** Department of Clinical Pharmacology, Faculty of Medical Sciences, Kyushu University, Maidashi 3-1-1, Higashi-ku, Fukuoka 812-8582, Japan (T.Y., T.S.); Clinical Research Center, Fukuoka Mirai Hospital, Kashiiteriha 3-5-1, Higashi-ku, Fukuoka 813-0017, Japan (T.Y.); Department of Social and Behavioral Sciences, Harvard T.H. Chan School of Public Health, 677 Huntington Avenue, 7th Floor, Boston, Massachusetts 02115, USA (M.Z.); Department of Public Health, Graduate School of Medicine, The University of Tokyo, 7-3-1 Hongo, Bunkyo-ku, Tokyo 113-0033, Japan (M.Z.); Kubota Maternity Clinic, Fukuoka, Japan (S.K.); Kubota Life Science Laboratory Co., Ltd., Saga, Japan (S.K.); Department of Preventive Medicine and Public Health, Faculty of Medicine, Fukuoka University, Nanakuma 8-19-1, Jonan-ku, Fukuoka 814-0180, Japan (H.A.)

**Keywords:** walking, water, pool, pregnancy-induced hypertension, renal blood flow, urine volume, prevention, renin-angiotensin-aldosterone-aldosterone system

## Abstract

**Background:** This study aimed to examine the effect of pool walking on renal function in pregnant women.

**Methods:** Fifteen pregnant women (mean gestational age, 37.8 weeks) walked in a pool (depth 1.3 m) for 1 h. A few days later, they walked on a street for 1 h. Within each activity, the starting and ending levels of plasma renin activity (PRA) and serum aldosterone (SA) were compared using paired t-test. Total urine volume, creatinine clearance, and change in PRA levels between each activity were compared by t-test. Regression coefficients for total urine volume and creatinine clearance during pool walking were estimated by linear regression and additionally controlled for the change in PRA levels. Land walking served as the reference group.

**Results:** Within each activity, the renin-angiotensin-aldosterone levels were suppressed during pool walking: the mean starting and ending values of PRA and SA were 6.8 vs. 5.5 ng/mL/h (p=0.002) and 654 vs. 473 pg/mL (p=0.02), respectively. Compared to land walking, the decrease in PRA level was more evident in pool walking (−1.27 vs. 0.81 ng/mL/h, p=0.004), resulting in higher total urine volume and creatinine clearance in pool walking (both p<0.05). In regression analysis, after controlling for the change in PRA levels, the significantly elevated regression coefficients for total urine volume and creatinine clearance in pool walking were attenuated.

**Conclusions:** Pool walking may temporarily improve renal function in pregnant women, partly through the suppressed renin-angiotensin-aldosterone system.

**Clinical Trial Registration:** URL: https://upload.umin.ac.jp/cgi-bin/ctr/ctr_view_reg.cgi?recptno=R000010618

Unique Identifier: UMIN000009051

## INTRODUCTION

Preeclampsia, along with other hypertensive disorders of pregnancy including eclampsia, chronic (preexisting) hypertension, preeclampsia superimposed on chronic hypertension, and gestational hypertension, is the leading cause for maternal mortality (e.g., stroke and pulmonary edema) and the risk for adverse fetal outcomes (e.g., preterm birth) worldwide.^1,2,3,4^ Currently, up to 10% of pregnant women have hypertensive disorders of pregnancy,^5,6^ and the prevalence of preeclampsia, the major contributor for maternal death, has been increasing over time particularly in (but not limited to) developed countries such as the USA.^2,3^ The increased risk for preeclampsia may be attributable to the pandemic of obesity, along with predisposing metabolic disorders (hypertension and diabetes) and the delay in childbearing to the age when these disorders are more common.^2,3^

Although some studies investigated the pharmaceutical approach for the prevention of preeclampsia (e.g., antihypertensive and antiplatelet agents), the efficacy and safety of these agents are inconclusive and still inconsistent.^1,2,3,6,7,8,9^ Moreover, behavioral approaches, including maternal prenatal physical activities, may have a potential for preventing preeclampsia. However, advantages of behavioral interventions are also inconsistent, and currently, maternal physical activities during pregnancy are not a guideline recommendation.^1,6^ For example, in recent systematic reviews, prenatal maternal exercise was safe, had benefits for the fetus, and was not associated with neonatal complications and adverse childhood outcomes,^10^ whereas behavioral interventions (including the broad range of education, diet, exercise, and self-monitoring of blood glucose) did not prevent hypertensive disorders of pregnancy.^11^ To improve guidelines in maternal physical activities during pregnancy, focusing “specific” physical activities in the biological pathogenesis of preeclampsia should be a priority.

Although the biological pathogenesis of preeclampsia is widely uncertain, it is recognized not merely as a disease of high blood pressure/renal dysfunction but also as a systemic disease stemming from the placental hypoperfusion/hypoxia with oxidative stress, inflammation, and endothelial/angiogenesis modifiers.^1,2,6,12^ Additionally, recent studies elucidated a potential biological pathway via the renin-angiotensin-aldosterone system.^13,14,15,16^ In a mouse model overexpressing human angiotensinogen and renin, the suppressed renin-angiotensin-aldosterone system appeared to play a key role in preventing preeclampsia.^13,14^ In the same model, physical exercise improved preeclampsia risk features of circulating and placental soluble fms-like tyrosine kinase-1.^14^

In this context and in order to prevent preeclampsia, we wanted to further evaluate a specific prenatal physical activity that may improve renal/placental circulatory perfusion via suppressed renin-angiotensin-aldosterone system. In physiological research for aquatic activities, hydrostatic pressure, which increases autotransfusion at capillary arteries, appeared to improve circulatory perfusion and suppress the renin-angiotensin-aldosterone system.^17^ However, few studies have assessed the effect of maternal physical activities during pregnancy in water.^18,19^

Accordingly, the aim of the present study was to examine the effect of head-out pool walking, a modified aquatic activity of head-out water immersion,^17^ on renal function in pregnant women. Using clinical and renal function data, we determined whether pool walking improved renal function (urine production and creatinine clearance) in combination with suppressed renin-angiotensin-aldosterone system.

## MATERIALS AND METHODS

### Study settings

The data that support the findings of this study are not publicly available. If any person wishes to verify our data, they are most welcome to contact the corresponding author. This study was approved by the Kyushu University Hospital Clinical Research Ethics Board, Fukuoka, Japan (Approval number: 24045) and registered in the UMIN Clinical Trial Registration (Registration number UMIN000009051). Written informed consent was obtained from all patients. All procedures performed in this study involving human participants were in accordance with the ethical standards of the institutional research committee and with the 1964 Helsinki declaration and its later amendments or comparable ethical standards.

An indoor swimming facility of the Fukuoka Swimming Club located in Fukuoka, Japan was used for the prenatal aquatic activity of pool walking. Aquatic activity instructors, accredited for cardiopulmonary resuscitation, overviewed and led the pool walking activity. The Kubota Maternity Clinic in Fukuoka, Japan, a general obstetric and gynecological hospital that closed in 2017,^21^ conducted the pool walking program along with the Fukuoka Swimming Club from 2003 to 2017. In our previous empirical experience, pregnant women tended to improve ill-health conditions in the placenta such as venous distension and edema through pool walking (Figure 1^20^). Pool walking is common in Japan as an aquatic exercise, and indoor pool facilities usually provide a “walking” lane.

**Figure 1.**
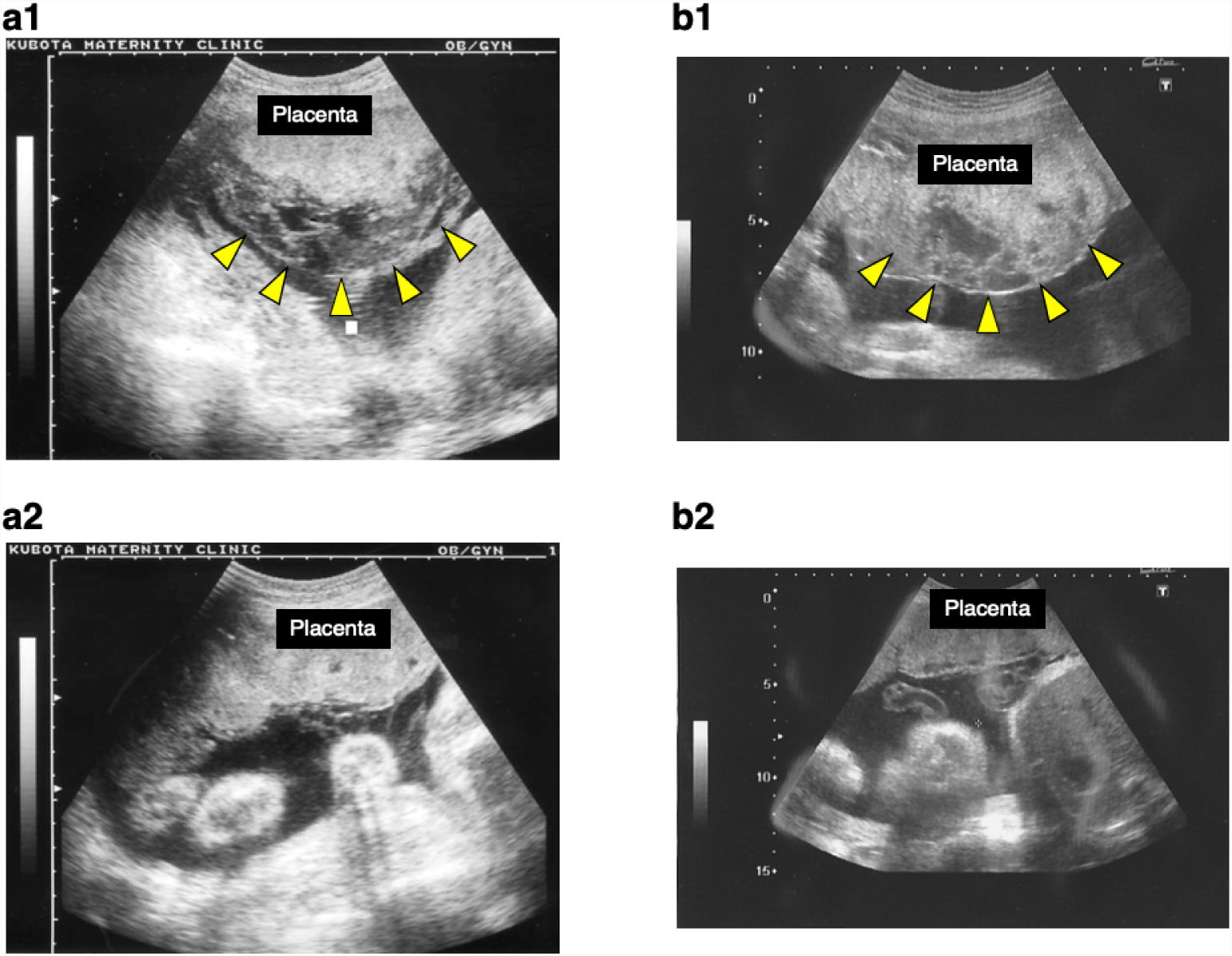
Empirical cases with improved venous distension and edema in the placenta by pool walking activity. Figures were adapted and modified from Reference 1 (Kubota 2017). In Case 1, (a1) the placental venous distension at the gestation age of 16 weeks (arrows) was improved at the gestation age of 30 weeks (a2). In Case 2, (b1) the placental edema (arrows) at the gestation age of 30 weeks was improved at the gestation age of 36 weeks (b2).

### Study design and participants

In this study, participants walked in the indoor pool with the head-out position for 1 hour at their own pace. The dimensions of the pool having six lanes were as follows: length 25 m (27.3 yds), depth 1.3 m (4.3 ft). The water temperature was set at 31-32°C, and the ambient room temperature at 29-30°C. A few days later (2 to 9 days), the same participants walked on a street, i.e., a conventional prenatal activity, near the Kubota Maternity Clinic for 1 hour at their own pace. We first planned to conduct this study in the typical cross-over design by randomizing half of the participants to the reverse sequence. However, due to a potential risk based on our empirical experience that land walking might induce labor,^20^ we finally decided not to choose the typical cross-over design to prioritize the safety of our participants and their babies. Therefore, every participant firstly did pool walking and then land walking. The weather conditions during land walking were fine/cloudy, the temperature was ∼23 to 26°C, and the humidity was ∼ 40 to 50%. During each activity, the participants were allowed to drink water freely. Dates for each activity were determined according to the schedule of the participants.

The eligible participants in this study were pregnant women (aged 20 to 39) with a normal pregnancy (gestational age, 37 weeks and above), who had prenatal care at the Kubota Maternity Clinic and had already participated in the pool walking program before the study period. Participants were ineligible if they presented cardiovascular disease, diabetes, hypertensive disorders of pregnancy, placenta previa/low-lying placenta, amniorrhexis, or anemia (hemoglobin <10 g/dl), or those considered not suitable for this study by physicians. We initially recruited 16 participants (11 participants in October 2012 and five participants in October 2013). One participant (37 years of age, 38.4 weeks of gestational age, and 35 sessions of pool walking before the study) was excluded due to delivery before land walking exercise; therefore, only 15 participants were included in the analysis. Since no previous studies with the effect of pool walking in pregnant women were available, sample size was calculated based on a previous study of water immersion^18^ and our empirical experience.^20^

### Assessments of clinical and renal function parameters

Body weight, systolic and diastolic blood pressures, and pulse rate were measured at the start and end of each walking activity. Blood samples were collected at the start and end of each walking activity, and the starting and ending levels of plasma renin activity (PRA), serum aldosterone (SA), hematocrit, and serum creatinine were measured.^18,19^ Within each walking activity, we calculated the changes in each parameter, subtracting the starting levels from the ending levels. According to previous studies, we defined the change in PRA as a potential indicator for the regulation of the renin-angiotensin-aldosterone system.^13,14,18,19^

Before each walking activity, all participants urinated. During each walking activity, all urine samples were collected, and the total amount of urine was recorded. With the total urine sample at the end of each walking activity, urine creatinine was measured. All laboratory measurements were performed at the Clinical Laboratory Center of Fukuoka City Medical Association, Fukuoka, Japan. We calculated creatinine clearance as follows: creatinine clearance = urine creatinine (mg/dL) × urine volume (mL/min)/[(serum creatinine at the start + serum creatinine at the end)/2 (mg/dL)].

### Statistical analysis

Between each activity, the starting levels of clinical and renal function parameters, as well as the changes in these parameters, were compared by t-test. The total urine volume and creatinine clearance were also compared by t-test. Within each activity, the starting and ending levels of each parameter were compared by paired t-test. The analyses for total urine volume and creatinine clearance were restricted to 14 participants since it was not possible to collect a sample from one participant during land walking.

We estimated the regression coefficient (β) and 95% confidence interval (CI) of pool walking for total urine volume and creatinine clearance, respectively. The land walking activity served as a reference. Age, body mass index, and gestational age were adjusted as confounding variables (model 1). In model 2, the decrease of PRA was additionally controlled as a potential mediating variable, i.e., this variable would not confound the association between walking activity and the renal function but might instead help to *explain* the observed difference between activities. No other potential covariates (systolic and diastolic blood pressures, pulse rate, and hematocrit) were included due to the small sample size with non-significant results in prior analyses (data not shown).

Alpha value was set at 0.05, and all p-values were two-sided. Data were analyzed using JMP pro 12 (SAS Institute, Cary, NC, USA) and STATA/MP 13.1 (Stata-Corp, College Station, TX, USA).

## RESULTS

The baseline characteristics of the participants are shown in Table 1. The starting levels of each parameter did not differ between pool and land walking activities (Table 1). The mean amount of water intake did not differ between activities (mean ± SD: 179 ± 76 versus 185 ± 56 mL, p=0.81).

**Table 1.** Baseline characteristics of study participants.

| Characteristics | Mean (SD) |  | p<sup>*</sup> |
| --- | --- | --- | --- |
|  | Before pool walking | Before land walking |  |
|  | (n = 15) | (n = 15) |  |
| Age, y | 32 (4) | - | - |
| Height, cm | 160.1 (5.4) | - | - |
| Body weight before pregnancy, kg | 51.8 (6.5) | - | - |
| Body mass index before pregnancy, kg/m<sup>2</sup> | 20.2 (2.3) | - | - |
| Prior attendance to walking in water, time | 25 (13) | - | - |
| Gestational age, week | 37.8 (1.0) | 38.2 (1.0) | 0.34 |
| Body-weight, kg | 60.4 (6.8) | 60.2 (6.7) | 0.94 |
| Body mass index, kg/m<sup>2</sup> | 23.5 (2.2) | 23.5 (2.1) | 0.93 |
| Systolic blood pressure, mmHg | 115 (12) | 111 (13) | 0.49 |
| Diastolic blood pressure, mmHg | 69 (7) | 68 (9) | 0.91 |
| Pulse rate, beat per minute | 81 (11) | 79 (14) | 0.63 |
| Hematocrit, % | 35.2 (2.1) | 34.9 (2.0) | 0.66 |
| Plasma renin activity, ng/mL/h | 6.8 (3.4) | 6.3 (4.0) | 0.71 |
| Serum aldosterone, pg/mL | 654 (518) | 674 (407) | 0.91 |

<sup>\*</sup> t-test.

Within each activity, the starting and ending levels of clinical and renal function parameters are shown in Table 2. During pool walking, the ending levels of body-weight, systolic and diastolic blood pressures, and hematocrit did not differ from the starting levels. However, the ending levels of pulse rate, PRA, and SA were significantly decreased (Table 2 and Figure 2). The mean starting and ending levels of PRA were 6.8 ± 3.4 versus 5.5 ± 2.9 ng/mL/h (p=0.002), respectively; the mean starting and ending levels of SA were 654 ± 518 versus 473 ± 304 pg/mL (p =0.02), respectively. By contrast, during land walking, the starting and ending levels of the examined parameters did not differ (although a tendency of decrease in body weight and pulse rate was observed, Table 2).

**Table 2.**
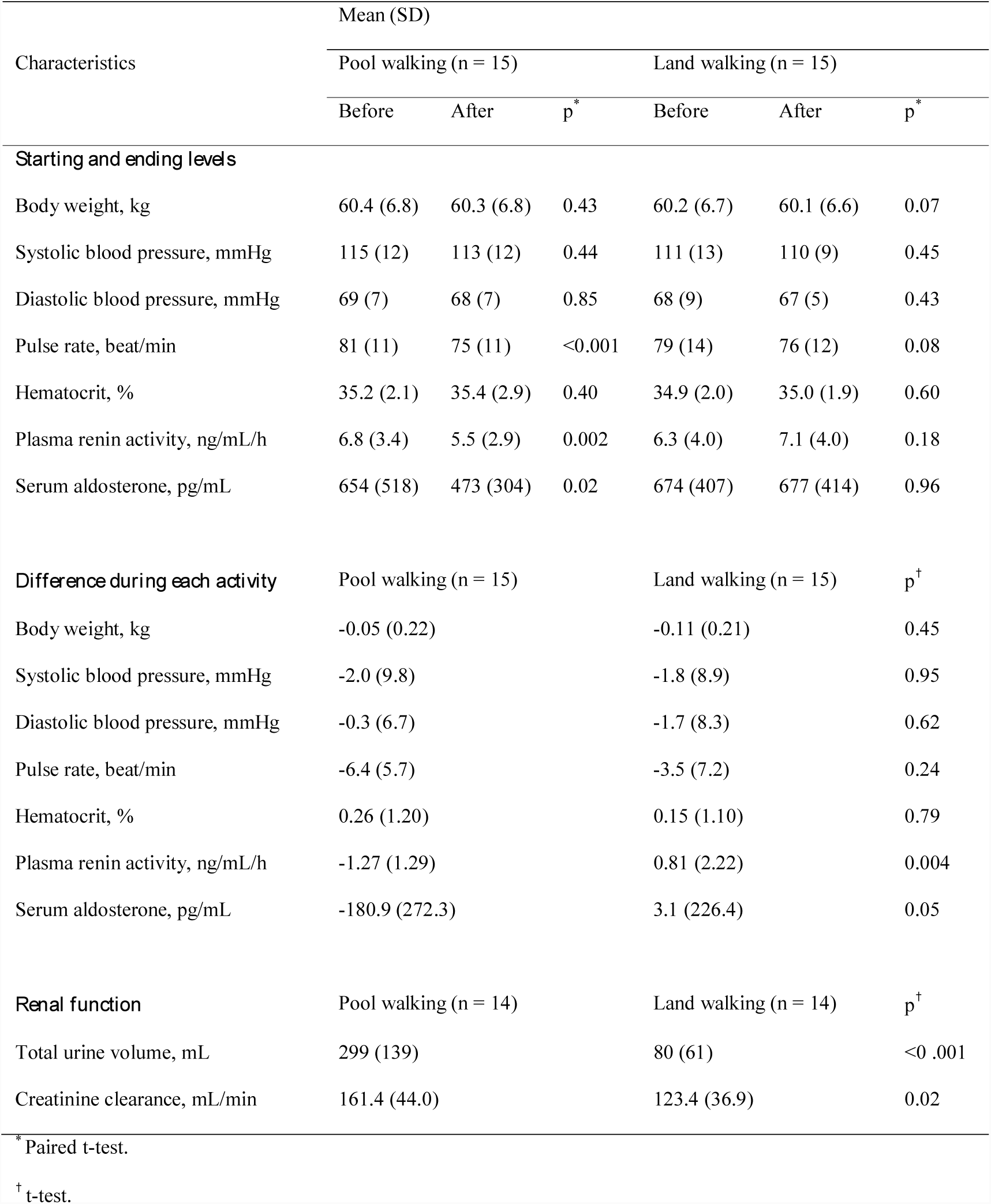
Starting and ending levels of clinical and renal function parameters within each activity.

**Figure 2.**
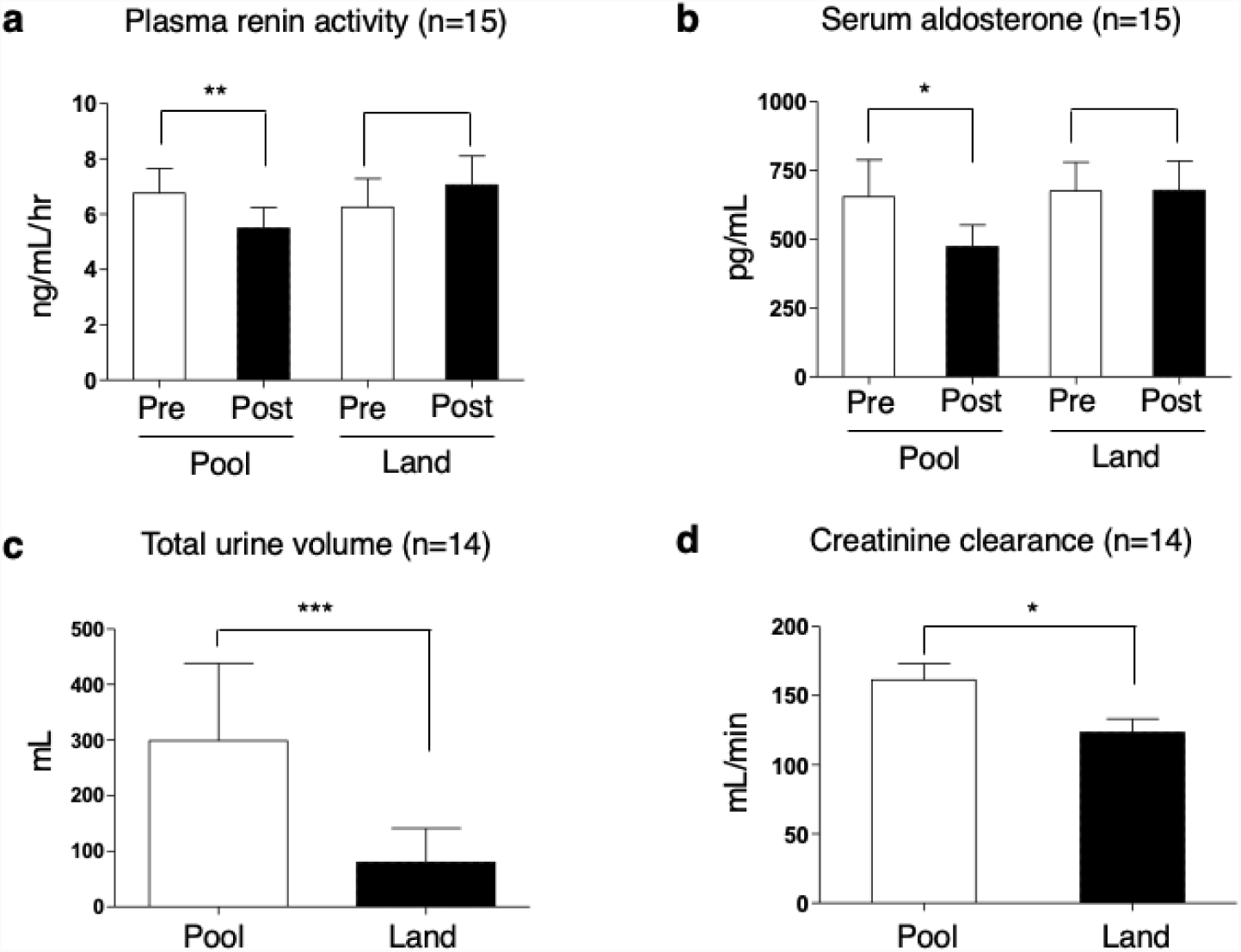
Changes of renal function parameters during 1-hour pool and land walking activities among 15 pregnant women. (a) Change of plasma renin activity level, (b) change of serum aldosterone level, (c) total urine volume, and (d) creatinine clearance. P-values are for (a, b) paired t-test and (c, d) t-test. For total urine volume and creatinine clearance, data were available for 14 participants. * p<0.05; ** p<0.01; *** p<0.001.

The results for the between-activity comparisons are shown in Table 2. Compared to land walking, the changes in body weight, systolic and diastolic blood pressure, pulse rate, and hematocrit did not differ in pool walking. However, the change in PRA significantly differed (Table 2) between pool and land walking activities being −1.27 ± 1.20 vs. 0.81 ± 2.22 ng/mL/h (p=0.004), respectively. Moreover, the change in SA differed marginally between pool and land walking activities (Table 2). As a result, compared to land walking, total urine volume and creatinine clearance were significantly higher in pool walking (Table 2 and Figure 2) being the mean total urine volume in pool and land walking activities 299 ± 139 vs. 80 ± 61 mL (p<0.001), and the mean creatine clearance in pool and land walking activities 161.4 ± 44.0 vs. 123.4 ± 36.9 mL/min (p=0.02), respectively.

The regression coefficients for total urine volume and creatinine clearance were significantly elevated in pool walking compared with land walking (model 1, Table 3), β_pool walking_ = 209 (95% CI, 123, 296) for total urine volume and β_pool walking_ = 32.2 (95% CI, 1.70, 62.7) for creatinine clearance. However, after controlling for the change in PRA levels, the elevated regression coefficients for total urine volume and creatinine clearance in pool walking were attenuated (model 2, Table 3), suggesting that the renal function was partly improved by pool walking through the suppressed renin-angiotensin-aldosterone levels.

**Table 3.**
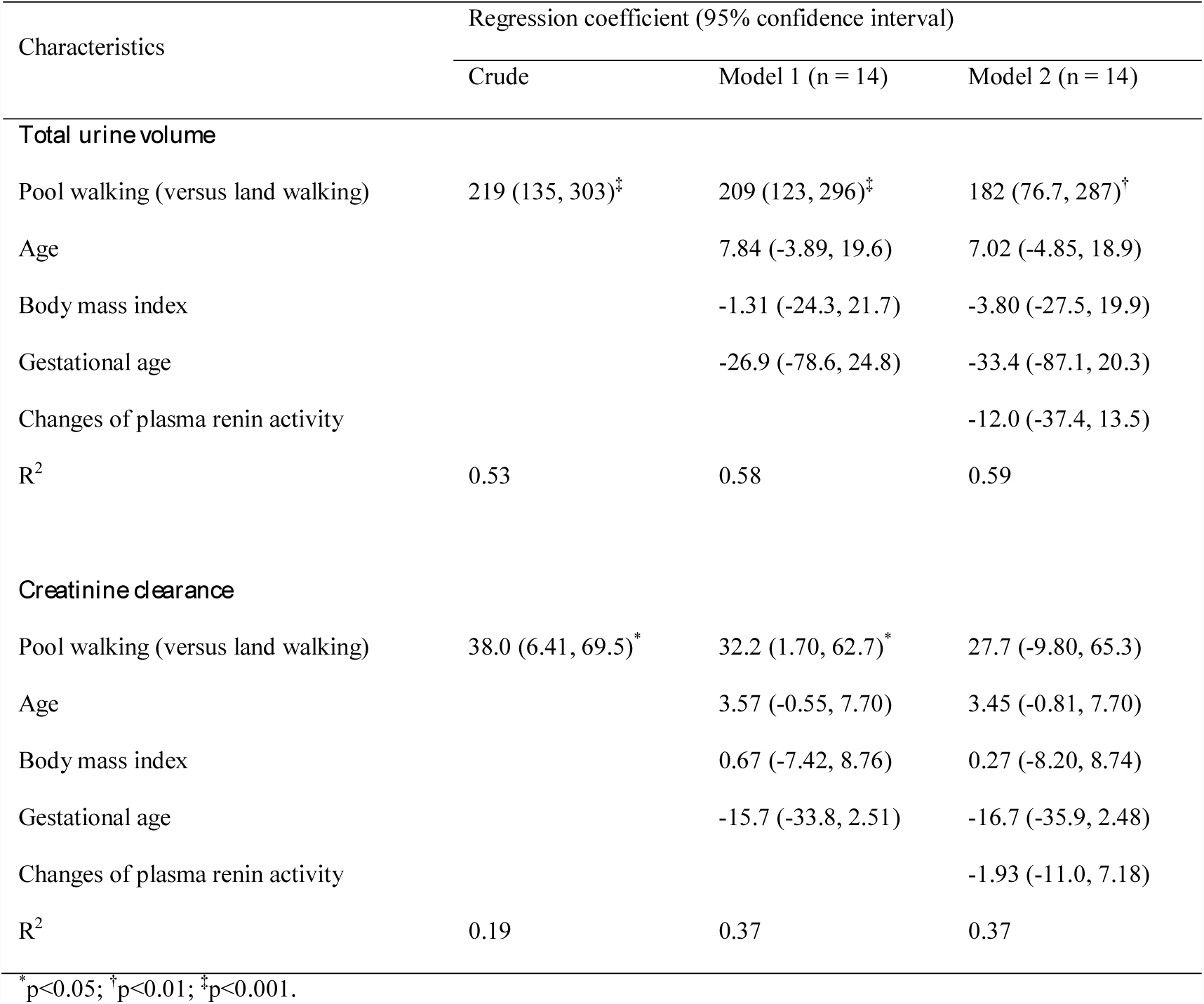
Regression coefficients of pool walking for total urine volume and creatine clearance estimated by linear regression.

## DISCUSSION

This study was the first to find that prenatal aquatic activity of head-out pool walking appeared to improve renal function, in combination with the regulation of the renin-angiotensin-aldosterone system. Compared to a conventional prenatal activity of land walking, pool walking suppressed the PRA and SA levels, resulting in improved urine production and creatinine clearance. Our findings suggested that pool walking might improve circulatory perfusion, including placental/renal perfusion, through hydrostatic pressure and that suppressed renin-angiotensin-aldosterone levels might result in an enhanced renal function. Our results also supported previous findings of head-out aquatic activities^17^ and potential biological pathogenesis from a mouse model.^13,14^

Hydrostatic pressure might be mainly attributable to the advantage of our pool walking activity. Previous studies suggested that hydrostatic pressure increased perfusion and autotransfusion at capillary vessels, particularly in the lower extremities, resulting in increased cardiac preload/output and decreased systemic vascular resistance.^17,18,22,23^ In addition, the renin-angiotensin-aldosterone system and sympathetic activity appeared to be suppressed in head-out water immersion with improved systemic/renal perfusion.^17,23,24^ In our study, we observed increased urine volume and creatinine clearance, suppressed renin-angiotensin-aldosterone levels, and decreased pulse rate during pool walking, which indeed suggests improved circulatory perfusion via hydrostatic pressure. Another advantage of pool walking might be buoyancy for pregnant women. On land, the enlarged pregnant uterus might compress the inferior vena cava and abdominal veins through the force of gravity, leading to decreased cardiac preload/output.^25,26^ By contrast, buoyancy in water might release the gravity-induced compression and improve cardiac preload/output.^17^ Conventional physical activities such as land walking might have some benefits,^1,6,10^ whereas exercise on land may have potential risks to reduce renal perfusion and urinary volume through vasoconstriction.^17^

Our pool walking activity had a unique feature of light-intensity physical activity with the head-out, vertical posture in water.^17^ Therefore, this prenatal activity was safe (“walk” as normal walking on land), had advantages of hydrostatic pressure, and required no special skills (swimming/breathing techniques). By contrast, the efficacy of other types of aquatic activities without the head-out position such as swimming might be inconsistent. In a previous study, moderate to strong swimming stimulated the renin-angiotensin-aldosterone system and increased PRA and catecholamine levels.^27^ In our empirical experience, compared to swimming, pool walking tended to be more comfortable for pregnant women since they preferred pool walking (the attendance rates for swimming and pool walking activities were ∼20% and ∼80%, respectively).^20^ In addition, pool walking should be more “fun” compared to water immersion, in which participants just stood/sat still in the water for several hours.^18^

Our study had some limitations. First, in order to prioritize the safety of our participants and their babies, we were not able to conduct our study in the typical cross-over design, and every participant first walked in water and then walked on land. This one-arm design might potentially carry-over the effect of pool walking to land walking. However, our participants walked on land at least two days after pool walking, and we confirmed that the baseline characteristics did not differ between the two activities. In addition, previous studies stated that PRA levels recover to baseline levels within a few hours after aquatic activities.^27^ Therefore, this limitation might not affect our sconclusions. Second, in addition to our small sample size, patients with hypertensive disorders in pregnancy were excluded, thereby limiting the generalizability of the results. In addition, other circulatory parameters (arterial diameters and arterial blood flow)^18,28^ and hormonal pathways (atrial natriuretic peptide)^24^ were not available. However, our empirical experience might support improved placental perfusion by pool walking (Figure 1).^20^ Third, although our data demonstrated temporal changes in the renin-angiotensin-aldosterone levels and renal function, our study did not assess the long-term and dose-response effects. Indeed, we did not observe the between-activity differences in the baseline characteristics, which might suggest that only one session of pool walking activity may not prevent preeclampsia risk along with other hypertensive disorders of pregnancy. Therefore, future studies are needed to clarify the long-term effect of pool walking with a randomized control design and to explore the potential benefits on cardiovascular health among not only pregnant women but also those with lifestyle-related diseases such as obesity.

In conclusion, this novel, pool walking activity may temporarily improve renal function by suppressing the renin-angiotensin-aldosterone system in pregnant women. Aquatic facilities are widely available at community-level (e.g., public indoor pools, as well as swimming pools in school, hotel, and fitness club) at least in developed countries such as USA and Japan. Hence, stakeholders including researchers, healthcare providers, and policymakers should focus further on the benefits of aquatic physical activities with hydrostatic pressure to tackle the norm of hypertensive disorders of pregnancy.

## Acknowledgement

We are grateful to Yuichi Takahashi and Hikari Yasuda in the Fukuoka Swimming Club for helping us to conduct this study and Professor Ichiro Kawachi in the Harvard T.H. Chan School of Public Health for his insightful inputs on this study. We would like to thank Editage (www.editage.jp) for English language editing.

## Author Contributions

Tatsuya Yoshihara: Conceptualization, resources, formal analysis, writing - original draft, supervision, and writing - review and editing. Masayoshi Zaitsu: Formal analysis, writing - original draft, and writing - review and editing. Shiro Kubota: Conceptualization, resources, supervision, and writing - review and editing. Ichiro Kawachi: Writing - review and editing. Hisatomi Arima: Formal analysis and writing - review and editing. Toshiyuki Sasaguri: Conceptualization, supervision, and writing - review and editing.

## References

1. American College of Obstetricians and Gynecologists, Task Force on Hypertension in Pregnancy. Hypertension in pregnancy. Report of the American College of Obstetricians and Gynecologists’ Task Force on Hypertension in Pregnancy. Obstet Gynecol. 2013;122:1122–1131.

2. Steegers EA, von Dadelszen P, Duvekot JJ, Pijnenborg R. Preeclampsia. Lancet. 2010;376:631–644.

3. Seely EW, Ecker J. Clinical practice. Chronic hypertension in pregnancy. N Engl J Med. 2011;365:439–446.

4. Schlembach D, Homuth V, Dechend R. Treating Hypertension in Pregnancy. Curr Hypertens Rep. 2015;17:63.

5. Vest AR, Cho LS. Hypertension in pregnancy. Curr Atheroscler Rep. 2014;16:395.

6. Sava RI, March KL, Pepine CJ. Hypertension in pregnancy: Taking cues from pathophysiology for clinical practice. Clin Cardiol. 2018;41:220–227.

7. Askie LM, Duley L, Henderson-Smart DJ, Stewart LA, PARIS Collaborative Group. Antiplatelet agents for prevention of preeclampsia: a meta-analysis of individual patient data. Lancet. 2007;369:1791–1798.

8. Banhidy F, Acs N, Puho EH, Czeizel AE. The efficacy of antihypertensive treatment in pregnant women with chronic and gestational hypertension: a population-based study. Hypertens Res. 2010;33:460–466.

9. Dodd JM, Louise J, Deussen AR, Grivell RM, Dekker G, McPhee AJ, Hague W. Effect of metformin in addition to dietary and lifestyle advice for pregnant women who are overweight or obese: the GRoW randomised, double-blind, placebo-controlled trial. Lancet Diabetes Endocrinol. 2019;7:15–24.

10. Davenport MH, Meah VL, Ruchat SM, Davies GA, Skow RJ, Barrowman N, Adamo KB, Poitras VJ, Gray CE, Jaramillo Garcia A, Sobierajski F, Riske L, James M, Kathol AJ, Nuspl M, Marchand AA, Nagpal TS, Slater LG, Weeks A, Barakat R, Mottola MF. Impact of prenatal exercise on neonatal and childhood outcomes: a systematic review and meta-analysis. Br J Sports Med. 2018;52:1386–1396.

11. Brown J, Alwan NA, West J, Brown S, McKinlay CJ, Farrar D, Crowther CA. Lifestyle interventions for the treatment of women with gestational diabetes. Cochrane Database Syst Rev. 2017;5:CD011970.

12. El-Sayed AAF. Preeclampsia: A review of the pathogenesis and possible management strategies based on its pathophysiological derangements. Taiwan J Obstet Gynecol. 2017;56:593–598.

13. Falcao S, Stoyanova E, Cloutier G, Maurice RL, Gutkowska J, Lavoie JL. Mice overexpressing both human angiotensinogen and human renin as a model of superimposed preeclampsia on chronic hypertension. Hypertension. 2009;54:1401–1407.

14. Genest DS, Falcao S, Michel C, Kajla S, Germano MF, Lacasse AA, Vaillancourt C, Gutkowska J, Lavoie JL. Novel role of the renin-angiotensin-aldosterone system in preeclampsia superimposed on chronic hypertension and the effects of exercise in a mouse model. Hypertension. 2013;62:1055–1061.

15. Jebbink J, Wolters A, Fernando F, Afink G, van der Post J, Ris-Stalpers C. Molecular genetics of preeclampsia and HELLP syndrome - a review. Biochim Biophys Acta. 2012;1822:1960–1969.

16. Mol BWJ, Roberts CT, Thangaratinam S, Magee LA, de Groot CJM, Hofmeyr GJ. Preeclampsia. Lancet. 2016;387:999–1011.

17. Pendergast DR, Moon RE, Krasney JJ, Held HE, Zamparo P. Human Physiology in an Aquatic Environment. Compr Physiol. 2015;5:1705–1750.

18. Elvan-Taşpinar A, Franx A, Delprat CC, Bruinse HW, Koomans HA. Water immersion in preeclampsia. Am J Obstet Gynecol. 2006;195:1590–1595.

19. Malha L, Sison CP, Helseth G, Sealey JE, August P. Renin-Angiotensin-Aldosterone Profiles in Pregnant Women with Chronic Hypertension. Hypertension. 2018;72:417–424.

20. Kubota S. Ninpu to akachan ni mananda hiesho to necchusho no kagaku. Tokyo, Japan: Tokyotoshoshuppan; 2017. [in Japanese]

21. Zaitsu M, Yoshihara T, Nakai H, Kubota S. Optimal Thermal Control with Sufficient Nutrition May Reduce the Incidence of Neonatal Jaundice by Preventing Body-Weight Loss Among Non-Low Birth Weight Infants Not Admitted to Neonatal Intensive Care Unit. Neonatology. 2018;114:348–354.

22. Epstein, M. Renal effects of head-out water immersion in man: implications for an understanding of volume homeostasis. Physiological Reviews. 1978;58(3),529–581.

23. Lazar JM, Khanna N, Chesler R, Salciccioli L. Swimming and the heart. Int J Cardiol. 2013;168:19–26.

24. Shiraishi M, Schou M, Gybel M, Christensen NJ, Norsk P. Comparison of acute cardiovascular responses to water immersion and head-down tilt in humans. J Appl Physiol. 2002;92:264–268.

25. Lanni SM, Tillinghast J, Silver HM. Hemodynamic changes and baroreflex gain in the supine hypotensive syndrome. Am J Obstet Gynecol. 2002;187:1636–1641.

26. Kember AJ, Scott HM, O’Brien LM, Borazjani A, Butler MB, Wells JH, Isaac A, Chu K, Coleman J, Morrison DL. Modifying maternal sleep position in the third trimester of pregnancy with positional therapy: a randomised pilot trial. BMJ Open 2018;8:e020256. doi:10.1136/bmjopen-2017-020256.

27. Vigas M, Celko J, Juránková E, Jezová D, Kvetnanský R. Plasma catecholamines and renin activity in wrestlers following vigorous swimming. Physiol Res. 1998;47:191–195.

28. Carter HH, Spence AL, Ainslie PN, Pugh CJA, Naylor LH, Green DJ. Differential impact of water immersion on arterial blood flow and shear stress in the carotid and brachial arteries of humans. Physiol Rep. 2017;5:e13285.

